# Connecting self-report and instrumental behavior during incubation of food craving in humans

**DOI:** 10.1101/2023.09.18.558282

**Authors:** Nicholas A Ruiz, Devlin Eckardt, Lisa A Briand, Mathieu Wimmer, Vishnu P Murty

## Abstract

Incubation of craving is a phenomenon describing the intensification of craving for a reward over extended periods of abstinence from reinforcement. Animal models employ instrumental markers of craving to reward cues to examine incubation, while homologous human paradigms often rely on subjective self-reports. Here, we characterize a novel human paradigm that showed strong positive relationships between self-reports and instrumental markers of craving. Further, both measures expressed non-linear relationships with time since last consumption, which parallels homologous animal paradigms of incubation of craving.

---

Experiencing craving, or a strong desire to use drugs, has recently been added as a diagnostic criterion for substance use disorders (SUDs) in the DSM-5. Importantly, craving has been shown to play a significant role in relapse (Vafaie & Kober, 2022). Thus, it is critical to form a comprehensive understanding of craving to develop novel therapeutic treatments for SUDs. A hallmark of addiction is the incubation of craving, where craving for a reward increases over extended periods of abstinence. Initially reported by Gawin & Kleber, 1986, abstinent cocaine users’ self-reported craving escalated over weeks to months when exposed to drug-related cues (i.e., a needle), which had downstream consequences on the probability of relapse. Since this report, the incubation of craving has been replicated in humans and has founded one of the most prominent animal models of addiction across drugs (Neisewander et al., 2000, Grimm et al., 2001, Pickens et al., 2011, Li et al., 2015) and natural rewards (Grimm, 2020).

In animal models of incubation of craving, rodents learn to perform a behavior (e.g., lever press) that is paired with a cue to earn a reward. Craving is later measured by quantifying behavioral responses to the cue in the absence of reinforcement, during extended periods of abstinence. In these models, craving reliably displays an inverted U-shaped relationship across time, such that it is highest at moderate stages of abstinence (Shalev et al., 2001, Grimm et al., 2003). While these animal models have provided a foundation for understanding the mechanisms of craving and relapse (Pickens et al., 2011, Loweth et al., 2014, Dong et al., 2017, Reiner et al., 2019, Altshuler et al., 2020, Venniro et al., 2021), the translational value of rodent models into humans has been limited based on differences in how craving is probed across species (Liu et al., 2023).

Human studies of incubation have mainly relied on self-report measures of craving which represent hedonic feelings toward the prospect of reward rather than motivational vigor evoked by the drug (Nava et al., 2008, Bedi et al., 2011, Wang et al., 2013, Parvaz et al., 2016, Li et al., 2016) or food cues (Coutinho et al., 2018). In animal models, craving is typically queried by lever presses reinforced by cues previously associated with reward, which is thought to more directly represent motivational vigor (Liu et al., 2023). This contrast creates a disconnect within the literature where animal models of incubation assess craving by measuring the behaviors animals perform to seek rewards and human models measure the associated feelings. While in many cases these features of reward-seeking are intertwined, in cases of substance use they could become disassociated. Indeed, the utility of subjective measures of craving in predicting reward-seeking behavior has been questioned in the past (Kranzler et al., 1999, McKay 1999, Perkins, 2009). Therefore, empirical evidence is needed to bridge the gap between the methods used to study craving in rodent and human models of incubation.

The goal of the current project is to validate a novel paradigm in humans that integrates behaviors akin to rodent assays of craving (i.e., instrumental responses) with the subjective human experience (i.e., self-report) in the space of natural rewards: palatable foods. We chose to characterize food craving as an entry into the domain due to its relative ease to assess. We predicted (1) a positive relationship between instrumental responses and subjective self-reports of craving and (2) that both instrumental responses and self-reports will show an inverted U-shaped relationship with time since last consumption.

The study hypotheses, analyses, and sample size were pre-registered prior to data collection (AsPredicted #107775). Participants were recruited via Prolific.ac (Palan & Schitter, 2018). 131 participants ranging in age between 18-35, with normal or corrected-to-normal vision who lived in the United States and spoke fluent English were recruited for study 1 with 103 meeting the exclusion criteria. Study 2 was a direct replication of study 1 with a sample size based on a power analysis of the linear mixed effects model predicting instrumental behavior by time and liking, which indicated an n = 150 was required for power = 0.80 with alpha = 0.05. We recruited 208 participants with 164 meeting the exclusion criteria. Participants in both studies were paid at a rate of $15/hour.

Informed consent and stimuli were presented using Inquisit (Grootswagers, 2020). Participants first completed a demographics questionnaire, reported their hunger level and the last time they ate. Participants first listed 10 of their favorite foods and 10 neutral foods they neither liked nor disliked to eat **(Fig 1a)**. For both lists, participants were told to be specific, providing specific brands, locations to attain the food, or the person who makes it. Participants were then shown each of the listed foods individually, in randomized order, and completed ‘pre-ratings’ **(Fig 1b)**. They were asked to rate how much they liked that item, how much they craved that item, and when they last consumed that item. Liking and craving were measured on a scale from 0 to 100, where 0 indicated “not at all” and 100 indicated “extremely”. For time since last consumption, participants responded on a scale from 0 to 100 that had 7 anchors that included “within 10 minutes”, “within 90 minutes, “within 16 hours”, “within 7 days”, “within 2 months”, “within 2 years”, and “greater than 2 years ago”. After completing the pre-ratings, participants completed a short distractor task.

**Figure 1.**
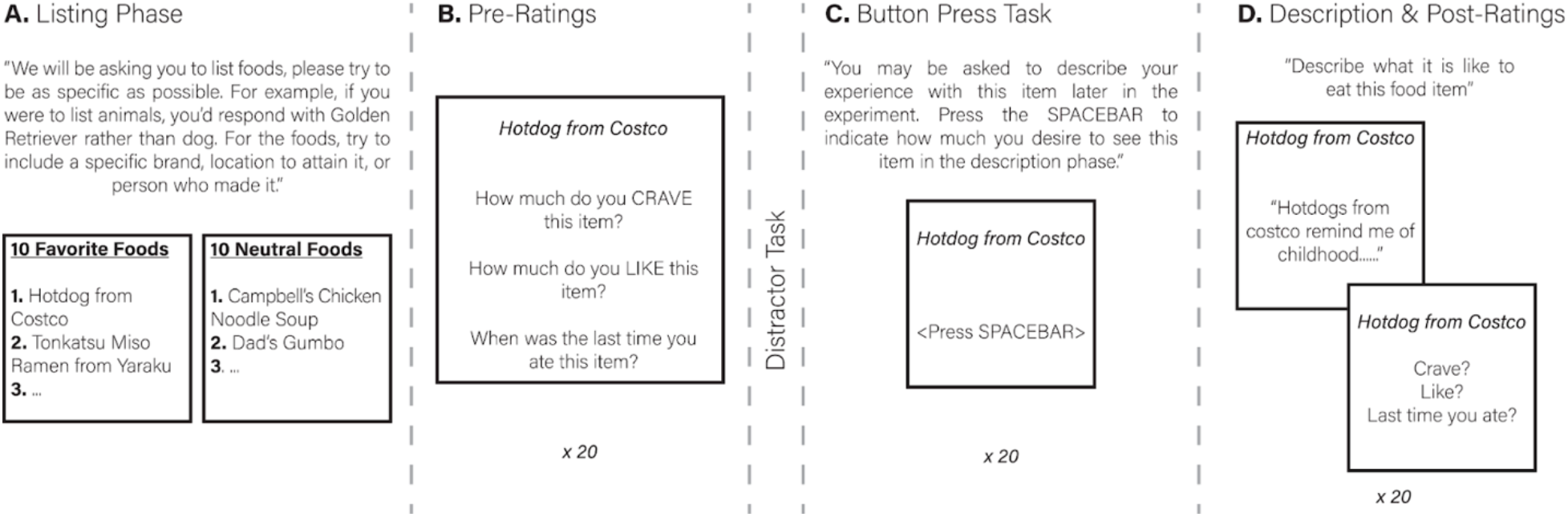
**A**. Listing phase. Participants were asked to list 10 of their favorite foods and 10 neutral foods, food items they neither liked nor disliked. They were instructed to be specific, including brand names, locations to attain the food, or the person who made the food. **B**. Pre-ratings. Participants rated each item on a scale from 0-100 indicating how much they craved the item, how much they liked the item, and the last time they consumed the item. **C**. Button press task. Participants were told they may be given the opportunity to describe an experience with each food item in a later phase of the experiment. They were instructed to press the spacebar to indicate how much they desired to see that item in the later description phase and that more spacebar presses indicated an increased desire to describe the item. **D**. Description and post-ratings. Participants were asked to write at least 5 sentences describing what it was like to eat each food item. Immediately following their description, they rated that item for craving, liking, and time since last consumption.

Participants next completed an instrumental behavior task **(Fig 1c)** and were told that for each food item, they would have an opportunity to describe an experience with that food item later in the experiment. As we were interested in using this task as a proxy to measure craving akin to rodent models, we told participants to press the spacebar to indicate their desire to provide an experience with the item, with more spacebar presses indicating greater desire. We selected instrumental responses to retrieve a memory of the food item to mirror the presentation of cues in the absence of reinforcement in rodent incubation paradigms. All 20 items were shown in a randomized order and participants could freely press the spacebar for 10s before moving onto the next item. Finally, participants were told to write at least five sentences describing what it is like to eat all of the food items **(Fig 1d)**. As the data from the description phase was beyond the scope of the current project, it was not analyzed and will be examined in a subsequent report (AsPredicted #107775).

Before data analysis, we excluded data on the following criteria: (1) not responding to more than 10% trials in any phase of the task, (2) responding with a repetitive response for more than half of the trials in any phase of the task, (3) participants whose liking ratings differed from their assignment to favorite versus neutral lists using paired t-tests of p < 0.05. To examine the relationship between our main variables of interest and our dependent variables (DV), we fit linear mixed-effects models and used a model comparison approach. A baseline model included only a single predictor and a main model added additional predictors that we hypothesized would have a meaningful effect on the DV above and beyond the initial predictor. All models were computed on a trial-wise level and included “subject” as a random effect. Model fits were assessed using AIC and BIC and we conducted chi-squared tests to compare the models (see “supplemental_table.pdf”). Liking ratings were included in each of the baseline models as the initial predictor to differentiate liking and craving (see “supplemental_preregistration_difference.pdf”).

We first examined the relationship between subjective craving and time since last consumption using the data from the pre-ratings **(Fig 2a)**. In study 1, we found that including linear and non-linear effects of time to our baseline model significantly increased model fit (baseline: craving ∼ liking + [1|subject]; winning model: craving ∼ time + I(time^2) + liking + [1|subject]; model comparison: χ^2^(2) = 13.37, p = 0.001). The winning model showed that craving was significantly predicted by liking (β(1968) = 0.91, p < 0.001, SE = 0.02, t = 58.85), time (β(1973) = 0.31, p = 0.006, SE = 0.11, t = 2.76), and quadratic time (β(1977) = -0.002, p = 0.03, SE = 0.0009, t = - 2.15). For study 2 **(Fig 3a)**, following the methodology of previous replication reports (Bryan et al., 2019), we report one-tailed hypothesis tests on the winning model from study 1. In study 2, we found craving was significantly predicted by liking (β(3148) = 0.89, p < 0.001, SE = 0.01, t = 73.42), time (β(3168) = -0.16, p = 0.046, SE = 0.10, t = -1.69) and quadratic time (β(3180) = 0.001, p = 0.03, SE = 0.0007, t = 1.90).

**Figure 2.**
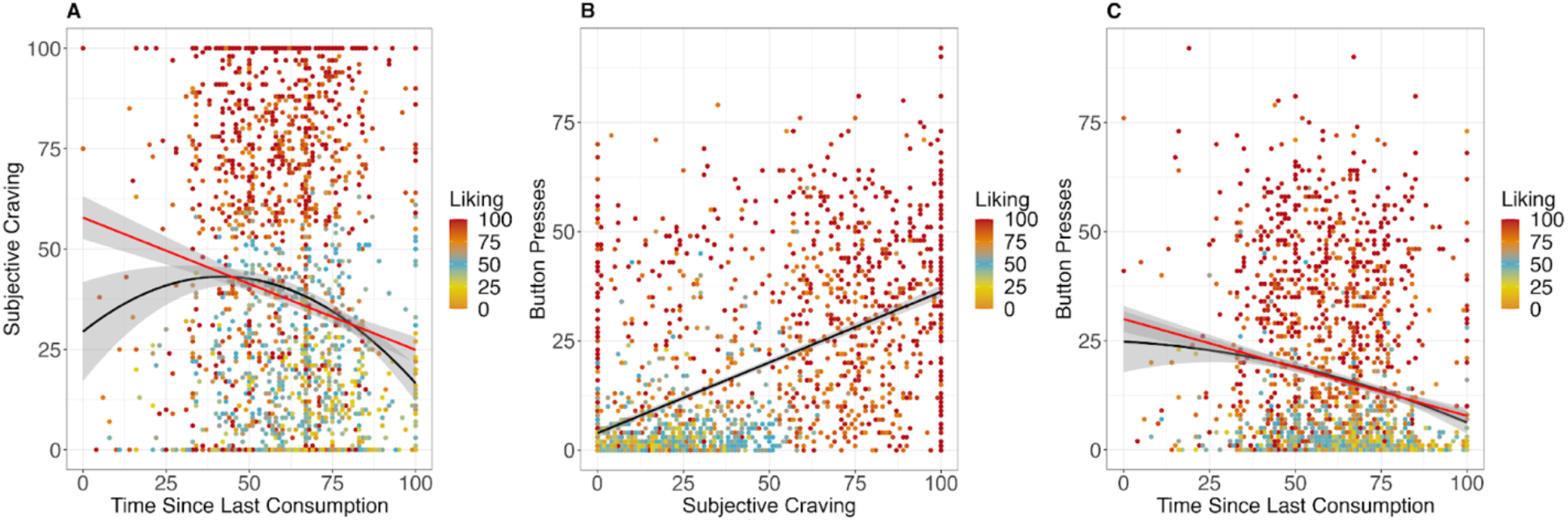
Results from Study 1. **A**. Subjective Craving by Time. Study 1 found a significant linear effect of time on subjective craving (red line, p=0.006). Further, there was a significant inverted U-shaped relationship between time since last consumption and craving (black line, p=0.03). **B**. The relationship between button presses and self-report of craving. Study 1 found a significant relationship between button presses and self-report of craving (p<0.001). **C**. Instrumental Behavior by Time. Study 1 found a significant linear relationship between time and instrumental behavior (red line, p=0.005). Further, there was a significant inverted U-shaped relationship between time since last consumption and instrumental behavior (black line, p=0.001), mirroring data from rodent incubation paradigms.

**Figure 3.**
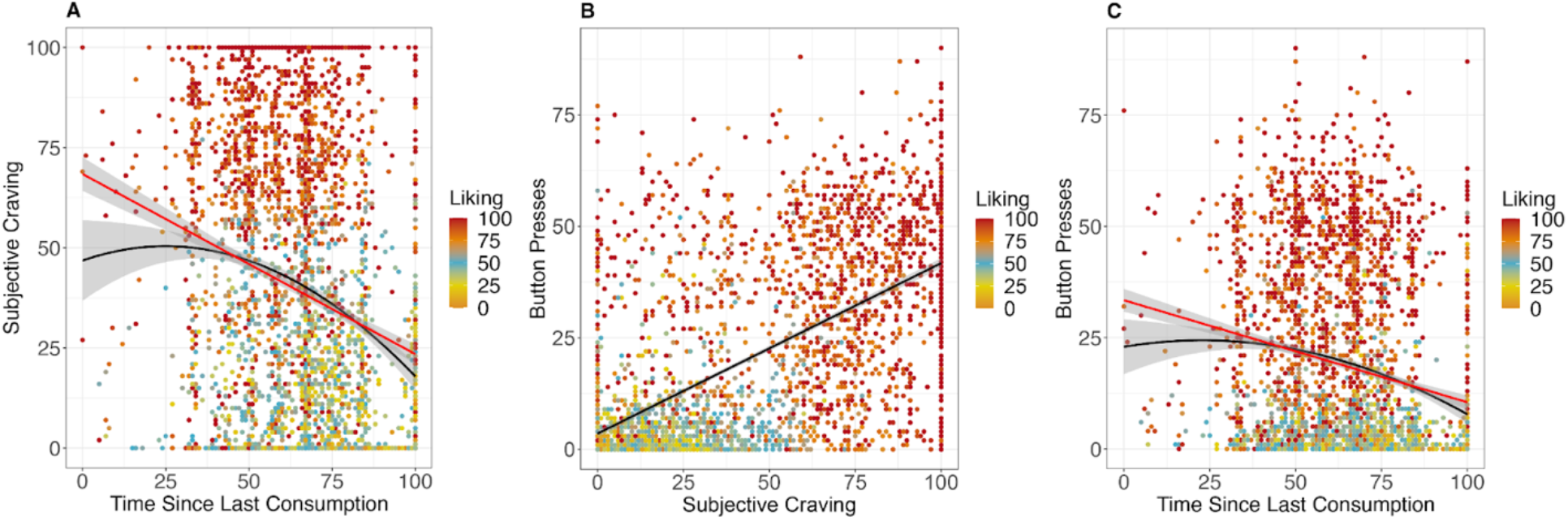
Results from Study 2. **A**. Subjective Craving by Time. Study 2 found a significant linear effect of time on subjective craving (red line, p=0.05, one-tailed). Further, there was a significant inverted U-shaped relationship between time since last consumption and craving (black line, p=0.005, one-tailed), replicating Study 1. **B**. The relationship between button presses and self-report of craving. Study 2 found a significant relationship between button presses and self-report of craving (p<0.001, one-tailed), replicating Study 1. **C**. Instrumental Behavior by Time. Study 2 found a trend towards linear relationship between time and instrumental behavior (red line, p=0.08, one-tailed). Further, there was a trend towards inverted U-shaped relationship between time since last consumption and instrumental behavior (black line, p=0.06, one-tailed).

We next explored the relationship between button presses and time since last consumption **(Fig 2c)**. In study 1, we found that including linear and non-linear effects of time to our baseline model when predicting button presses increased our model fit (model comparison: χ^2^(2) = 13.51, p = 0.001). The winning model showed that button presses were predicted by liking (β(1982) = 0.49, p < 0.001, SE = 0.001, t = 47.51), time (β(1991) = -0.21, p = 0.005, SE = 0.08, t = -2.80), and quadratic time (β(1999) = 0.002, p = 0.001, SE = 0.0006, t = 3.27). In study 2 **(Fig 3c)**, we again report one-tailed hypothesis tests of the winning model from study 1. Study 2 found that button presses were significantly predicted by liking (β(3170) = 0.51, p < 0.001, SE = 0.009, t = 59.08), but there was no significant effect of time (β(3200) = -0.94, p = 0.08, SE = 0.07, t = - 1.39) or quadratic time (β(3216) = 0.0008, p = 0.06, SE = 0.0005, t = 1.56). Given the conflicting results across both studies, we combined the data across both studies. In this combined sample, we find a significant effect of liking (β(5154) = 0.50, p < 0.001, two-tailed, SE = 0.007, t = 75.17), time (β(5193) = -0.14, p = 0.006, SE = 0.05, t = -2.75) and quadratic time (β(5217) = 0.001, p = 0.002, two-tailed SE = 0.0004, t = 3.16).

Finally, we explored the relationship between self-report of craving and button presses. In study 1 **(Fig 2b)**, we found that including self-reports of craving in our baseline model significantly increased our model fit when predicting button presses (baseline: presses ∼ liking + [1|subject]; winning model: presses ∼ craving + liking + [1|subject]; model comparison: χ^2^(1) = 89.48, p < 0.001). The winning model showed that button presses were predicted by both liking (β(2038) = 0.36, p < 0.001, SE = 0.02, t = 22.69) and self-reports of craving (β(1991) = 0.13, p < 0.001, SE = 0.01, t = 9.56). In study 2 **(Fig 3b)**, we again report one-tailed hypothesis tests of the winning model from study 1. Study 2 found that button presses were significantly predicted by liking (β(3264) = 0.34, p < 0.001, SE = 0.02, t = 26.09) and self-reports of craving (β(3201) = 0.19, p < 0.001, SE = 0.01, t = 16.24).

These findings bridge human and animal models of craving. First, we found that individuals reported higher subjective craving for items they had not consumed for moderate amounts of time compared to those consumed recently or in the far past. Critically, these findings were specific to cravings and not just general hedonic affect, as we controlled for liking in all of our analyses. This relationship dovetails with prior human incubation of drug craving (Nava et al., 2008, Bedi et al., 2011, Wang et al., 2013, Li et al., 2014, Parvaz et al., 2016, Coutinho et al., 2018) and extends the literature by replicating this effect in the context of a natural reward in a non-clinical population. By characterizing multiple behavioral assessments of craving in humans, the current results pave a path toward investigation into the heterogeneous systems involved in addiction across species (Kalivas et al., 2005).

Next, we showed a similar relationship between time since the last consumption and instrumental button presses to share a memory of the food reward, which parallels animal models of incubation that focus on motivated behavior rather than measures of affect. Critically, we found a positive relationship between subjective craving and instrumental behavior such that when participants reported higher subjective craving, they performed more button presses to recall details of a reward. Previous work has captured objective measures for craving in humans (Ooteman et al., 2006), using physiological (Peter et al., 2000, Liu et al., 2022) and neural markers (Parvaz et al., 2016, Koban et al., 2022), which are difficult to execute in large-scale samples. We extend this prior work by providing an instrumental marker of craving that is more homologous to animal models and can be implicated in large-scale behavioral studies.

One interesting feature of our paradigm, that differs from other assessments of incubation in both the rodent and human literature, is that we probed motivational vigor in response to the opportunity to retrieve a memory about rewards rather than a conditioned cue (i.e., showing a lighter to a nicotine addict). We believe this design feature more closely mirrors models of craving and relapse that are centered on retrieving prior experiences with a reward. In line with this decision, there is a growing body of work showing involvement of the hippocampus, a region critical for memory retrieval, in cue-evoked craving. The hippocampus has shown craving-related activation in response to cue-exposure in drug (Kilts et al., 2001, Schneider et al., 2001, Smolka et al., 2005), and natural rewards (Crockford et al., 2005, Pelchat et al., 2004, Stevenson & Francis, 2017). The exact role of the hippocampus in cue-induced craving is unclear but given its role in retrieval of autobiographical memories (Maguire, 2001, Sheldon & Levine, 2016), it is possible representations of previous drug use, or the rewards related to drug use, are brought online during cue-exposure. Due to the episodic nature of these memories (Tulving, 1985), the individual may feel as if they are re-living the previous drug using experience which may lead to drug-seeking behavior. Further, more recent research has shown that autobiographical memory retrieval, albeit not in the context of rewards, is associated with both hedonic and motivational value (Speer et al., 2014, Speer & Delgado, 2020).

The use of memory retrieval also provides a scaffold for a better understanding of why reward-related memories can remain strong over long periods of abstinence, resulting in relapse. Currently, theories of reward learning that serve as a foundation for drug addiction cannot account for the incubation phenomenon (Wise, 2004). Specifically, they predict reward memories should fade over time when there is no direct exposure or reinforcement with the reward or reward-related cues. However, the plethora of incubation data shows the opposite, craving for reward can in fact increase even without reward or cue exposure. Memory consolidation processes provide a possible mechanism by which memory for the reward or related cues can intensify over time (Buzsaki, 2015). Therefore, a dissection of these learning systems is required to fully understand how reward memories can incubate without re-exposure.

There remains an open question regarding the use of food and food craving in the current study. We chose to assess food craving due to its relative ease to assess. However, we predict the current results will extend beyond food craving and into the drug domain due to the similarity between incubation in drug and natural rewards (Grimm, 2020) as well as the overlapping neural circuitry concerning reward and pleasure between reward domains (Kragel et al., 2023).

Additionally, the translation of subjective craving into behaviors to seek reward is not entirely clear. While we were not able to directly assay this in the current study, there is mixed evidence regarding the correlation between subjective craving and relapse. Some research show a positive relationship between the two (Bottender & Soyka, 1995, Stohs et al., 2019) while other literature has not (Kranzler et al., 1999, Perkins, 2009). These discrepancies necessitate future research discerning what features underlie human craving responses, or how different contextual features influence craving response (McKay 1999, Secades-Villa and Fernandez-Hermida, 2003). However, using paradigms similar to ours, which include self-reports of craving and instrumental button presses, could provide more reliable assays of craving, which could be of use in clinical communities.

## Supporting information

supplement pre-registration

## Acknowledgements

We have no conflicts of interest to disclose. We would like to thank Dr. Emily Cowan for feedback on early versions of the manuscript. We also thank Virginia Ulichney for assistance with statistical analysis.

