## supplemental table 1 for "Connecting self-report and instrumental behavior during incubation of food craving in humans"

| study | model | AIC | BIC |
|---|---|---|---|
| 1 | craving ~ liking + [1\|subject] | 17981 | 18004 |
| 1 | craving ~ time + I(time^2) + liking + [1\|subject] | 17972 | 18006 |
| 2 | craving ~ liking + [1\|subject] | 28851 | 28876 |
| 2 | craving ~ time + I(time^2) + liking + [1\|subject] | 28851 | 28888 |
| 1 | presses ~ liking + [1\|subject] | 16230 | 16252 |
| 1 | presses ~ craving + liking + [1\|subject] | 16142 | 16171 |
| 2 | presses ~ liking + [1\|subject] | 26536 | 26561 |
| 2 | presses ~ craving + liking + [1\|subject] | 26284 | 26315 |
| 1 | presses ~ liking + [1\|subject] | 16230 | 16252 |
| 1 | presses ~ time + I(time^2) + liking + [1\|subject] | 16220 | 16254 |
| 2 | presses ~ liking + [1\|subject] | 26536 | 26561 |
| 2 | presses ~ time + I(time^2) + liking + [1\|subject] | 26537 | 26574 |

**AIC and BIC table.** This table provides the AIC and BIC numbers for each model of each main analysis for both studies 1 and 2. The ‘winning’ model provided the lowest AIC and BIC scores with the rest of the associated statistics provided in the manuscript.
