## supplement pre-registration for "Connecting self-report and instrumental behavior during incubation of food craving in humans"

**Difference from pre-registration**

In the pre-registration we noted controlling for condition by using a binomial variable that represented high (favorite) vs low (neutral) liking. We opted to use the liking ratings from the pre-rating phase, as this allowed for more specificity and flexibility within each model.
